# Congruent visual cues speed dynamic motor adaptation

**DOI:** 10.1101/2023.02.01.526660

**Authors:** Sae Franklin, Raz Leib, Michael Dimitriou, David W. Franklin

**Affiliations:** Neuromuscular Diagnostics, Department of Sport and Health Sciences, Technical University of Munich, Germany; Physiology Section, Department of Integrative Medical Biology, Umeå University, Sweden; Munich Institute of Robotics and Machine Intelligence (MIRMI), Technical University of Munich, Munich, Germany; Munich Data Science Institute (MDSI), Technical University of Munich, Munich, Germany

**Keywords:** motor control, force field adaptation, motor learning, internal model, motor memory, additional visual cues

## Abstract

Motor adaptation to novel dynamics occurs rapidly using sensed errors to update the current motor memory. This adaption is strongly driven by proprioceptive and visual signals, that indicate errors in the motor memory. Here we extend this previous work by investigating whether the presence of additional visual cues could increase the rate of motor adaptation, specifically when the visual motion cue is congruent with the dynamics. Six groups of participants performed reaching movements while grasping the handle of a robotic manipulandum while an additional cue (red object) was connected to the cursor. After a baseline, either a unidirectional (3 groups) or bidirectional (3 groups) velocity dependent force field was applied during the reach. For each group, the movement of the red object relative to the cursor was either congruent with the force field dynamics, incongruent with the force field dynamics or constant (fixed distance). Participants adapted more to the unidirectional force fields than to the bidirectional force field groups. However, across both force fields, groups in which the visual cues matched the type of force field (congruent visual cue) exhibited higher final adaptation level at the end of learning compared to either the control or incongruent conditions. In all groups, we observed an additional congruent cue assisted the formation of the motor memory of the external dynamics. We then demonstrate that a state estimation based model that integrates proprioceptive and visual information can successfully replicate the experimental data.

**New & Noteworthy:** We demonstrate that adaptation to novel dynamics is stronger when additional online visual cues that are congruent with the dynamics are presented during adaptation, compared to either a constant or incongruent visual cue. This effect was found regardless of whether a bidirectional or unidirectional velocity dependent force field was presented to the participants. We propose that this effect might arise through the inclusion of this additional visual cue information within the state estimation process.

## Introduction

Humans adapt to new environments and tasks by updating their motor memories to these changed conditions. Even fundamentally novel environments can be learned by the sensorimotor system through the incorporation of sensory information (Farshchiansadegh et al. 2015; Liu et al. 2011; Ranganathan et al. 2014). For example, visual or proprioceptive sensory feedback can convey information regarding possible errors or rewards that trigger changes to the motor plan. Although it is clear that the motor system uses information coming from these sensory channels to understand the altered environment, especially when they carry essential or unique information, it is still unknown how additional and perhaps redundant information might also affect adaptation of our motor memories.

It has long been clear that motor adaption to changes in task dynamics is strongly driven by proprioceptive input (DiZio and Lackner 2000). There are many studies in which adaptation relies on proprioceptive information without the need for additional visual feedback. Critically, visually impaired people can learn new dynamics (DiZio and Lackner 2000). Similarly, eliminating visual feedback while experiencing external forces did not affect the speed or amount of adaptation to these forces (Franklin et al. 2007; McKenna et al. 2017; Scheidt et al. 2005). We can also rapidly adapt our grip forces to objects’ weight (Lukos et al. 2007; Westling and Johansson 1984) or other generated load forces (Leib et al. 2015) without visual information regarding the object. In all of these cases, in which the sensorimotor system needs to rely on sensory inflow (reafference), motor out flow (dead reckoning) or combination of two, an estimate of the current state (e.g. position and velocity) of the hand must be generated (Wolpert et al. 1995).

While visual information is not essential for motor adaptation, when present it may have a significant role in the learning process. Previous studies showed the existence of motor adaptation without the need for visual information (DiZio and Lackner 2000; Franklin et al. 2007; Scheidt et al. 2005) while others showed that we can adapt our motor plans based solely on visual feedback. For example, it has been shown that participants moving in a force channel which eliminates any proprioceptive error can learn novel dynamics only with visual position error feedback (Melendez-Calderon et al. 2011). Similar adaptation was found in proprioceptive deafferented patients that use visual information to adapt to external forces (Lefumat et al. 2016; Sarlegna et al. 2010; Yousif et al. 2015). Moreover, these patients can use visual information to some extent in order to adapt their grip force to load forces using vision (Hermsdörfer et al. 2008; Miall et al. 2019). It can be claimed that visual information triggers a more conscious strategy of adaptation (Hwang et al. 2006) which works only in specific cases (Forano et al. 2021; Howard et al. 2013).

Adaptation requires accurate estimates of internal states. To produce these, it is likely that the nervous system makes favorable use of all sensory modalities at its disposal. Indeed, it is widely believed that internal representations normally rely on the statistically optimal integration of different types of sensory feedback (Bays and Wolpert 2007; Körding and Wolpert 2004). For reaching movements in particular, vision and proprioception are weighted differently based on their direction-dependent precision (van Beers et al. 2002). Moreover, in the context of implicit motor adaptation, visual feedback can even lead to an appropriate modulation of muscle proprioceptive feedback itself (Dimitriou 2016; 2018). In short, visual feedback can play an important role in motor adaptation.

While visual feedback is not essential for motor learning, there is some evidence that it can affect the learning rate. Vision, combined with proprioceptive information, can improve the estimation regarding changes in the environment. The upgraded estimation can allow for faster learning since we can predict the perturbation in following movements. For example, in tasks involving visual shift of the hand spatial position, participants learn the perturbation slower when the visual information regarding hand position was blurry compared to sharp (Burge et al. 2008; Wei and Koerding 2010). This example demonstrates that visual information affects the uncertainty that we have regarding the observed information and change our ability to modify the internal representation of the environment.

Here we further investigate the role of visual feedback on the sensorimotor control system during adaptation. In contrast with previous studies which tested the effect of arbitrary (Bernardi et al. 2013) or distorted visual cues (McKenna et al. 2017), we focused on the effect of providing additional, potentially meaningful, visual cues. We assessed whether the existence of additional visual cues could affect the speed of adaptation to different force field types (unidirectional and bidirectional). Specifically, we tested if a visual cue that is congruent or incongruent with the experienced forces affects the learning of novel force field dynamics.

## Materials and Methods

### Subjects

Sixty neurologically healthy, right handed (Oldfield 1971) human subjects (34 males and 26 females) took part in the experiment (mean age of 24.3 ± 5.5 years). All participating subjects were naïve as to the purpose of the study and gave their informed consent prior to participating. Each subject participated in one experimental session of approximately 45 minutes each and was randomly assigned to one of the six conditions. The institutional ethics committee at the University of Cambridge approved the study.

### Experimental Apparatus and Setup

Subjects were firmly strapped into an adjustable chair in front of a robotic rig (Fig 1A). They made reaching movements with their right arm in the horizontal plane at approximately 10 cm below their shoulder level while the forearm was supported against gravity with an air sled. Subjects grasped the handle of a vBOT robotic interface, which was used to generate the environmental dynamics. The vBOT manipulandum is a custom-built planar robotic interface that can measure the position of the handle and generates forces on the hand {Howard:2009dj}. A six-axis force transducer (ATI Nano 25; ATI Industrial Automation) measures the endpoint forces applied by the subject at the handle. The position of the vBOT handle was calculated from joint-position sensors (58SA; IED) on the motor axes. Position and force data were sampled at 1 kHz. Visual feedback was provided using a computer monitor mounted above the vBOT and projected into the plane of the movement via a mirror. This virtual reality system covers the manipulandum, arm and hand of the subject, which obscures the location of the arm from their view.

**Figure 1.**
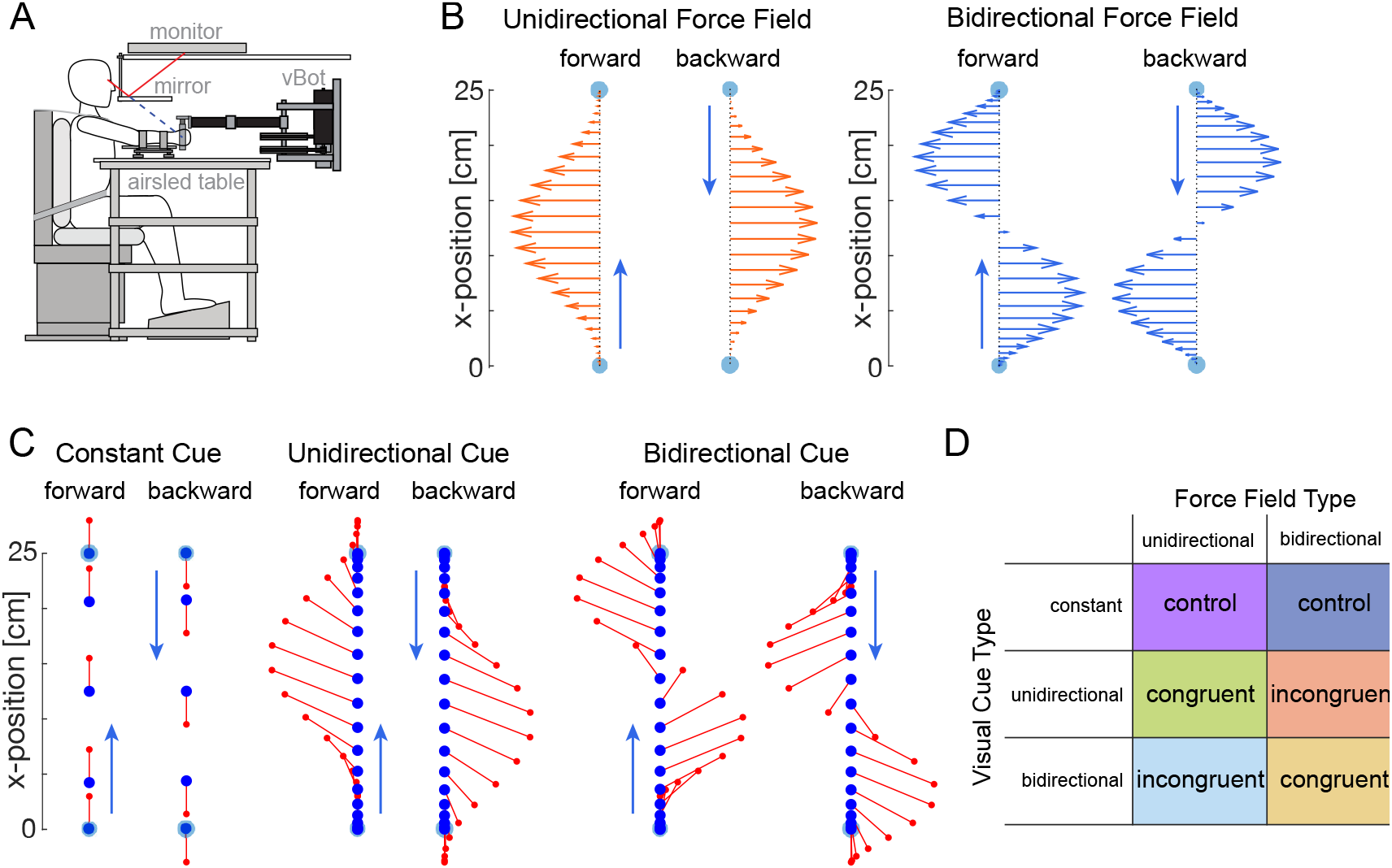
Experimental Setup. **A.** Participants were seated and grasped the handle of a robotic manipulandum with their right hand. Visual feedback about the task and the location of their hand was provided through a monitor reflected by a mirror such that it appeared in the plane of movement. The arm was supported by an airsled on a table. **B.** Two types of force fields were applied during an exposure phase. Three groups of participants performed movements within a unidirectional velocity dependent force field (left). Three other groups of participants performed movements in a bi-directional velocity dependent force field in which the sign of the forces switched halfway through the movement (right). **C.** Visual cues presented during movements depending on the group. Each of three groups for each force field was provided with a different visual cue during the movements. This visual cue was either constant (left), unidirectional (middle) or bi-directional (right). **D.** Six groups of participants performed the experiments with all possible combinations of visual cue type and force field type resulting in a control group, congruent group and incongruent group for each force field type.

### Experimental Paradigm

Subjects made alternating forward and backward point-to-point reaching movements between two targets. Trials were self-paced: the subject initiated each trial by moving the cursor (yellow circle of 0.5 cm radius representing the subject’s hand position) into the start circle (grey circle of 0.7 cm radius which became white once subjects had moved the cursor into the start circle). A small red circle (radius 0.3 cm) was visually attached to the end of the cursor via a thin red bar. This visual cue was located 2 cm directly in front of the cursor (in the direction of movement) when subjects were at rest but varied depending on the experimental condition. Once subjects had maintained the cursor within the start circle for 1s, a tone signalled to the subject to initiate a movement to a white target circle (radius 0.8 cm) located 25.0 cm away. The end of the movement was considered to have occurred once subjects had maintained the cursor within the target for 400 ms. Visual feedback was then provided about the success of the previous trial. A trial was considered successful if subjects did not overshoot the target, and the duration of the movement (between exiting the start and entering the target) was between 625 and 775ms. On successful trials the subjects received positive feedback (e.g. “good” or “great”) and a counter increased by one unit. Other messages were provided visually at the end of each trial to inform the subjects of their performance (either “too fast”, “too slow” or “overshot target”). All trials were analysed regardless of their success.

### Protocol

Subjects were randomly assigned to one of six groups (n=10). Subjects performed alternating forward and backward reaching movements in three phases of the experiment: pre-exposure (10 blocks), exposure (30 blocks), and post-exposure (5 blocks). Blocks consisted of 12 movements (6 forward and 6 backward reaches) such that each subject performed 540 movements total. These 12 movements consisted of 10 movements in the environmental dynamics (null field or force field) and 2 movements in a mechanical channel (one in each direction of movement), which were positioned randomly within the 12 trials. The mechanical channel (Milner and Franklin 2005; Scheidt et al. 2000) was implemented as a mechanical channel resisting lateral motion with a spring constant of 5,000 N/m and damping coefficient of 2 Ns/m. Both the pre-exposure and post-exposure phases were performed in the null field.

All groups experienced a velocity dependent force field during the exposure phase. This force field was a curl force field scaled by a value *k*, which varied throughout the movement as a function of the distance *d* between the start and target position:

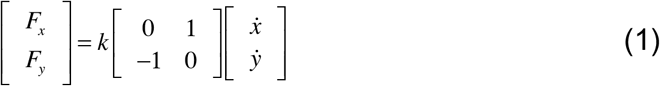

Three of the groups experienced a unidirectional velocity dependent force field during the exposure phase (Fig 1B, left). In the unidirectional force field, the value k varied as:

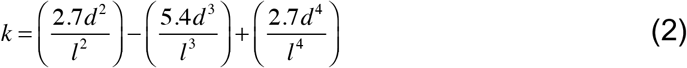

where 0 ≤ *d ≤l* and the length of movement *l* = 25.

The other three groups were presented with a bidirectional velocity dependent force field during the exposure phase (Fig 1B, right) where k was defined as:

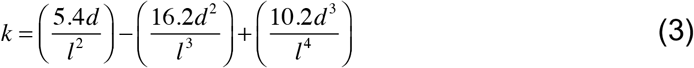

The direction of the force field was varied for half of the participants in each group by changing the sign of k.

Each group was presented with a visual cue throughout the experiment (Fig 1C). The visual cue was a red circle that was connected by a thin red line to the cursor. For one of the unidirectional field groups and one of the bidirectional field groups the location of the red circle was located x=0, y=±2 cm relative to the cursor (constant cue). That is, for forward movements it was located at y=+2 cm, whereas for backward movements it was located at y=-2 cm. For the other four groups (two unidirectional and two bidirectional), the visual cue was located x=k, y=±2 cm relative to the cursor, where k was defined using either equation 2 or 3. Specifically, for the two unidirectional force field groups, this visual cue could either be also unidirectional (congruent cue) or bidirectional (incongruent cue). Similarly, for the bidirectional force field the cue was either unidirectional (incongruent) or bi-directional (congruent). For all groups the value k was set to zero during the pre-exposure and post-exposure trials. This meant that during null field movements the visual cue was always located 2cm away from the cursor for all six groups. Overall, six groups of participants combined all possible combinations of visual cue type and force field type resulting in a control group, congruent group and incongruent group for each force field type (Fig 1D).

### Analysis

The data was analyzed using Matlab R2019a. Force and kinematic data was low-band pass filtered with a fifth-order, zero phase-lag Butterworth filter with a 40 Hz cutoff. Individual trials were aligned on movement onset. For each trial, we calculated measures of kinematic error or force compensation between 200 ms prior to leaving the start position until 200 ms after entering the target position.

For each non-channel trial, the absolute maximum perpendicular error (MPE) was calculated and used as a measure of the kinematic error. The MPE is the absolute maximum perpendicular distance between the hand trajectory and the straight line between the start and end targets.

To assess feedforward learning independent of co-contraction we analyzed the force compensation to the force field produced on channel trials (Smith et al. 2006). Force compensation is calculated by the regression between the force produced by participants into the wall of the simulated channel (lateral measured force) and the force needed to compensate exactly for the force field. Here the perfect compensatory endpoint force is determined on each trial using the forward velocity on each trial, and the magnitude of the force field k as calculated by equations 2 and 3. Here values in the null force field before learning (pre-exposure phase) should be close to zero.

We performed hypothesis-based planned comparisons and consider significance at the *p*<0.05 level for all statistical tests. ANOVAs were examined in JASP 0.16.4. For both the maximum perpendicular error and the force compensation values we performed an ANOVA with main factors of force field (2 levels: unidirectional and bidirectional) and condition (3 levels: constant, congruent and incongruent) on the final levels at the end of adaptation (final 10 blocks).

In order to examine differences in the time constant of adaptation and final asymptote we fit an exponential function to the force compensation for each trial throughout the adaptation phase of the experiment. To generate confidence intervals for our parameter estimates we performed block bootstrapping in which we left out each possible set of two participants from each group and fitted the remaining eight. We used the distribution of parameters across these 45 fits to estimate the confidence limits. To test whether each parameter varied between the two groups we generated all possible differences in each parameter from our bootstrap to generate a new bootstrap sample (45×45 samples). Differences in the magnitude and speed of learning were contrasted only across the same force fields.

### Model

To interpret these results, we suggest a framework in which the force field adaptation is based on estimating the scaling factor k using a state estimator. The state estimator integrates between proprioceptive and visual feedback in order to estimate the nature of the dependency of the scaling factor on the hand position while reducing the effect of any sensory noise. To simplify the problem, we suggest that the state estimator needs to estimate the coefficients of the polynomials describing the scaling factor. That is, for the unidirectional velocity dependent force field, the state estimator needs to estimate the values C_uni_=[2.7, −5.4, 2.7]^T^ and for the bidirectional velocity dependent force field the values C_bi_=[5.4, −16.2, 10.2]^T^. The observation, which is the input to the state estimator, includes proprioceptive and visual information about the scaling factor (depending on the experimental condition). This can be represented as:

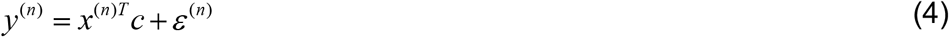

where y^(n)^ is the observations on the n-th trial, x^(n)^ is a vector representing the position dependency of the scaling factor, that is, 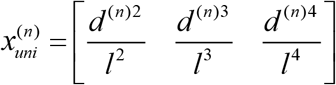 for the unidirectional case and 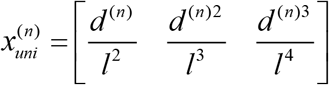 for the bidirectional case. ε^(n)^ represent random additive noise with normal distribution, *ε*^(*n*)^ *~ N*(0,*R*) where R represents the variance-covariance noise matrix. The R matrix was set as a diagonal matrix with values of 0.1 on the diagonal. Thus, we assume the same level of noise for both proprioceptive and visual sensory modalities. We implemented the state estimator as a Kalman filter (Kalman 1960). We initiate the system with an initial coefficients estimation, in this case, since the force field was turned on without any prior notice, we set the values to zero, 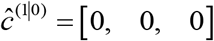 and noise added to proprioceptive or visual information as Gaussian noise with zero mean and 0.1 variance. Between experimental conditions we varied the initial uncertainty regarding the estimation. That is *p*^(1|0)^ was a diagonal matrix with different values between conditions. For the control condition and congruent condition we set the uncertainty values to 5 and 10 respectively for both the unidirectional and bidirectional force field. This means that for the congruent condition, since both the visual and proprioceptive modalities are providing the same information, the initial estimation of a null force field is more uncertain than for the control condition in which the information is providing only via a single sensory channel. In this case of higher uncertainty, the initial estimation is rapidly replaced with a more certain estimation. For the incongruent conditions we set the initial uncertainty to 1.5 and 1 for the unidirectional and bidirectional force fields respectively. In the case of incongruent visual information, we assume that the visual information is reducing the uncertainty regarding the initial estimation since the visual and proprioceptive information are not aligned, which requires gathering more information so to update the estimation. In this case the rate of adaptation will be slower than the learning rate in the congruent condition, or both the congruent and control conditions. To update the estimation, we used the Kalman algorithm

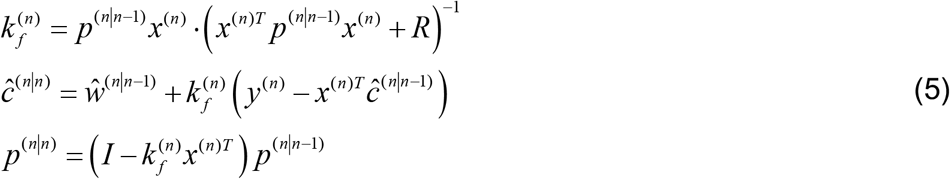

For each trial, the observation included 4 different points along the trajectory which were chosen randomly. Based on these four points we updated the coefficients estimation using the Kalman filter gains *k_f_*. Changing the number of points which are utilized in each trial affects the number of trials needed for the estimations to converge but not the nature of the estimation process. That is, reducing the number of points used in each trial will increase the number of trials needed and vice-a-versa. Based on the estimated coefficients, we estimated the scaling factor, 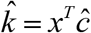, for each trial using 250 equally spaced points along the movement path and calculated mean square error between it and the scaling factor that was used in each experimental condition

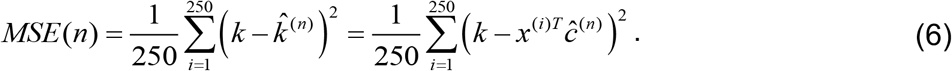

We run ten simulations for each condition same as the number of participants in each experimental group.

## Results

Six groups of participants performed alternating forward and backward discrete movements of the arm between two targets, with a small red circle (visual cue) connected to the cursor. After initial movements in a null field, either a unidirectional (3 groups) or bidirectional (3 groups) velocity dependent force field was applied during the movements. In each force field, the three groups were presented with the visual cue representing the force field (congruent cue), representing a different force field (incongruent cue) or unaffected (constant cue). Finally, the force field was removed (null field) and the visual cue was constant in all groups.

### Measures of Adaptation

In the pre-exposure phase, all six groups of participants performed relatively straight movements to the targets with little lateral kinematic error (Fig 2A,C). As participants were exposed to the force fields (exposure phase), the kinematic errors increased initially but reduced with further experience in the force fields. Several major effects are visible in the data. First there are clear differences in the reduction of kinematic error between the two force fields (unidirectional reduces error further than bidirectional). Second, for both force fields it appears that the congruent groups (yellow and green) learn slightly better (stronger reduction in MPE) than either the control (dark blue or purple) or incongruent (red and light blue) groups. The final level of MPE (last 10 blocks, Fig 2B,D)) was examined with an ANOVA with main factors of force field (2 levels) and condition (3 levels). There was a main effect of force field (F_1,54_=61.574; p<0.001; η^2^=0.5), but the main effect of condition failed to reach significance (F_2,54_=2.999; p=0.058; η^2^=0.049), and there was no interaction effect (F2,54=0.831; p=0.441; η^2^=0.013). Therefore, there is a strong effect of the force field on the reduction of the kinematic error during adaptation, but no clear evidence for differences across the three conditions.

**Figure 2.**
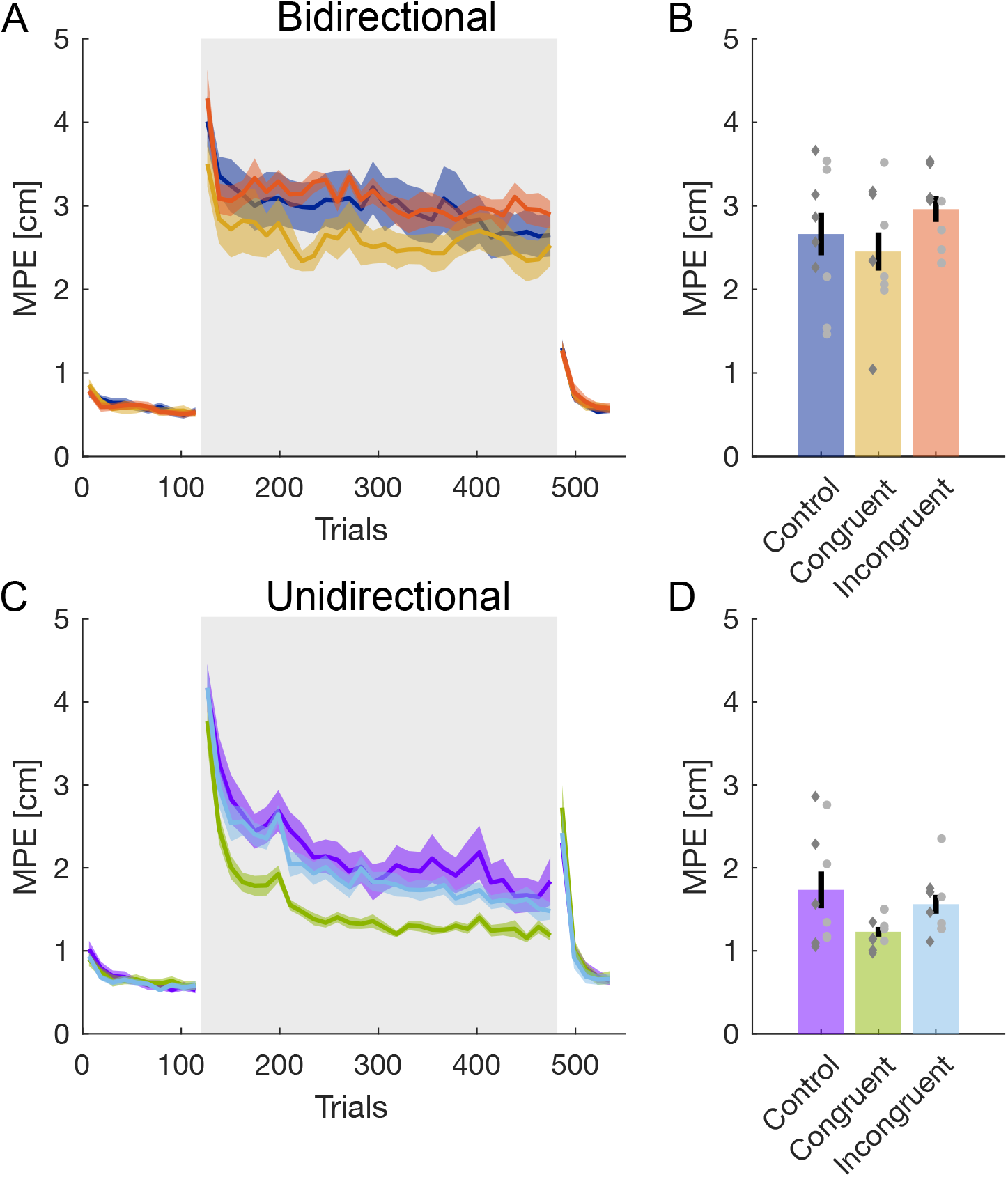
Kinematic error during adaptation. **A**. Absolute maximum perpendicular kinematic error throughout the experiment in the bidirectional force fields for the control (dark blue), congruent (yellow) and incongruent (red) groups. The mean (solid line) and standard deviation of the mean (shaded regions) across participants for each block in the experiments is shown. Shaded grey region indicates the period of force field exposure. **B.** Final levels of absolute maximum perpendicular error at the end of the exposure phase (last 10 blocks). Error bars are standard error of the mean, and individual points represent individual participants with the color indicating the direction of the force field (light grey circles: positive k; dark grey diamonds: negative k). **C.** Absolute maximum perpendicular kinematic error in the unidirectional force fields for the control (purple), congruent (green) and incongruent (light blue) groups. **D.** Final levels of absolute maximum perpendicular error at the end of the exposure phase (last 10 blocks).

To investigate the predictive compensation to the force fields independent of factors such as limb inertia or increases in co-contraction, we examined the force compensation throughout the experiment (Fig 3). During the initial pre-exposure phase the force compensation (relative to adapting to the subsequent force fields) remained close to zero as expected. Upon the start of the exposure phase, the force compensation for all groups increases, with a larger increase in the unidirectional force field groups (around 60%, Fig 3C) compared to the bidirectional force field groups (around 30%, Fig 3A). However, again we see that the two groups with the congruent visual cues have higher levels of force compensation at the end of adaptation that the other groups of participants. The final level of force compensation (last 10 blocks, Fig 3B, D) was examined with an ANOVA with main factors of force field (2 levels) and condition (3 levels). There were main effects of both force field (F_1,54_=191.968; p<0.001; η^2^=0.741) and condition (F_2,54_=5.491; p=0.007; η^2^=0.042), with no interaction effect (F_2,54_=1.107; p=0.338; η^2^=0.009). Therefore, there is a strong effect of the force field on the reduction of the kinematic error during adaptation, but also clear evidence for differences across the three conditions. A post-hoc comparison (Tukey) demonstrated that the congruent conditions had significantly higher force compensation than either the control (p=0.03) or incongruent conditions (p=0.009) but that there were no differences between the control and incongruent conditions (p=0.896) across the different force fields. Therefore, despite adapting to the same force fields and having the same visual presentation of the cursor representing hand position, participants that we also presented with an additional visual cue that matched the force field adapted more to the force fields (independent of the type of force field). That is, although all groups received visual feedback of the cursor motion (providing visual information regarding the errors induced by the force fields), additional congruent visual motion assists with the formation of the motor memory of the external dynamics.

**Figure 3.**
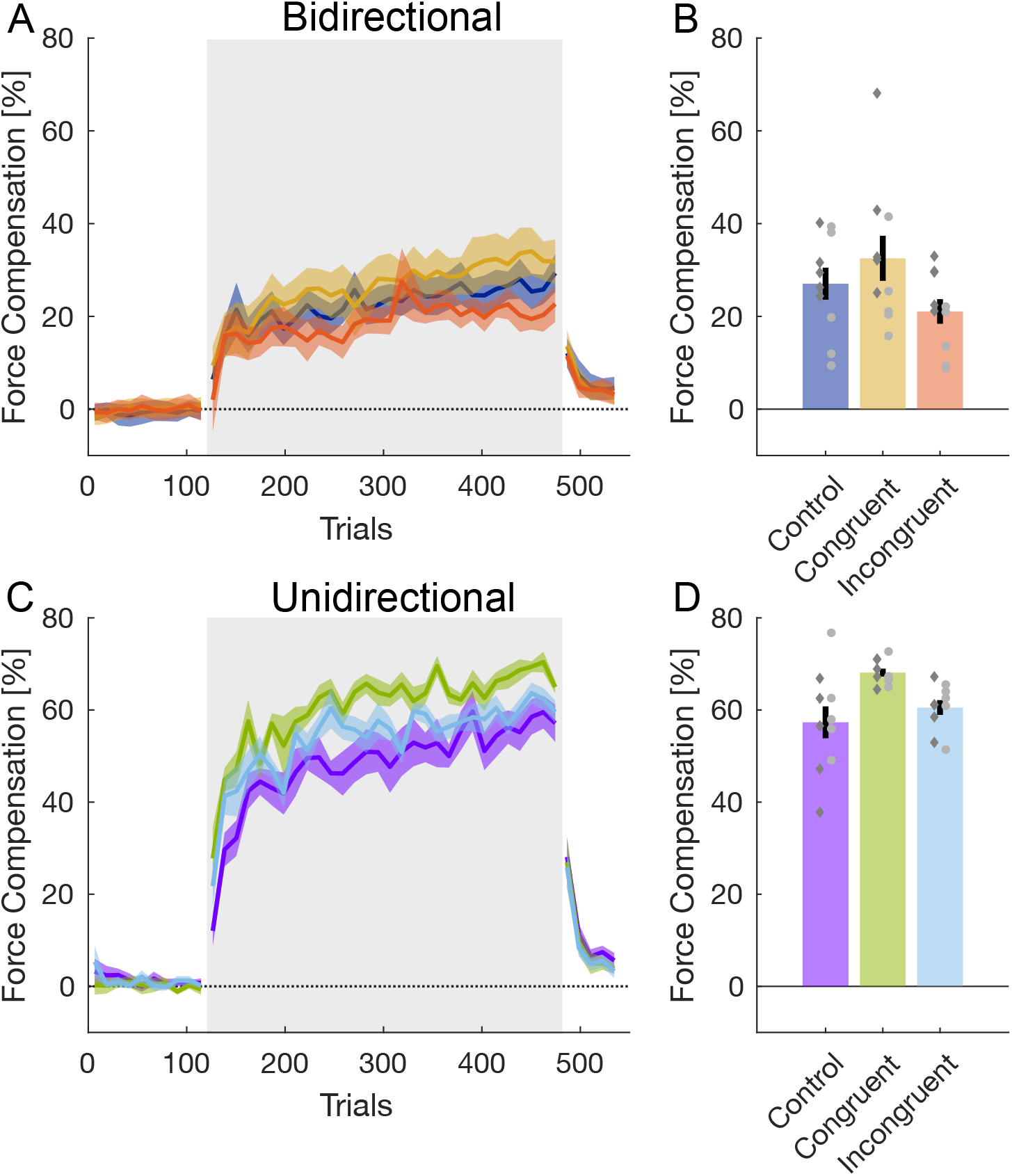
Predictive force compensation during adaptation. **A.** Force compensation as measured on channel trials throughout the experiment in the bidirectional force fields for the control (dark blue), congruent (yellow) and incongruent (red) groups. The mean (solid line) and standard deviation of the mean (shaded regions) across participants for each block in the experiments is shown. Shaded grey region indicates the period of force field exposure. **B.** Final levels of force compensation at the end of the exposure phase (last 10 blocks). Error bars are standard error of the mean, and individual points represent individual participants with the color indicating the direction of the force field (light grey circles: positive k; dark grey diamonds: negative k). **C.** Force compensation in the unidirectional force fields for the control (purple), congruent (green) and incongruent (light blue) groups. **D.** Final levels of force compensation at the end of the exposure phase (last 10 blocks).

### Learning Rates

The force compensation in the exposure phase was fit with an exponential function to determine the time contrast of adaptation and final asymptote levels for the six groups (Fig 4). For the bidirectional force field (Fig 4A-C), we find a higher final asymptote for the congruent compared to the control (p=0.044) and incongruent (p<0.001) groups, but also for the control compared to the incongruent group (p=0.0108). However, we find no differences in the time constant of adaptation across all three groups (all p>0.16). For the unidirectional force field (Fig 4D-F, we also find strong differences in the asymptote, with a larger adaptation for the congruent compared to the control (p<0.001) or incongruent groups (p<0.001), but here the incongruent group showed higher adaptation than the control group (p<0.001). The rates of adaptation to the unidirectional force field were higher for the congruent and incongruent groups than the control group (both p<0.001) but with no difference between them (p=0.436).

**Figure 4.**
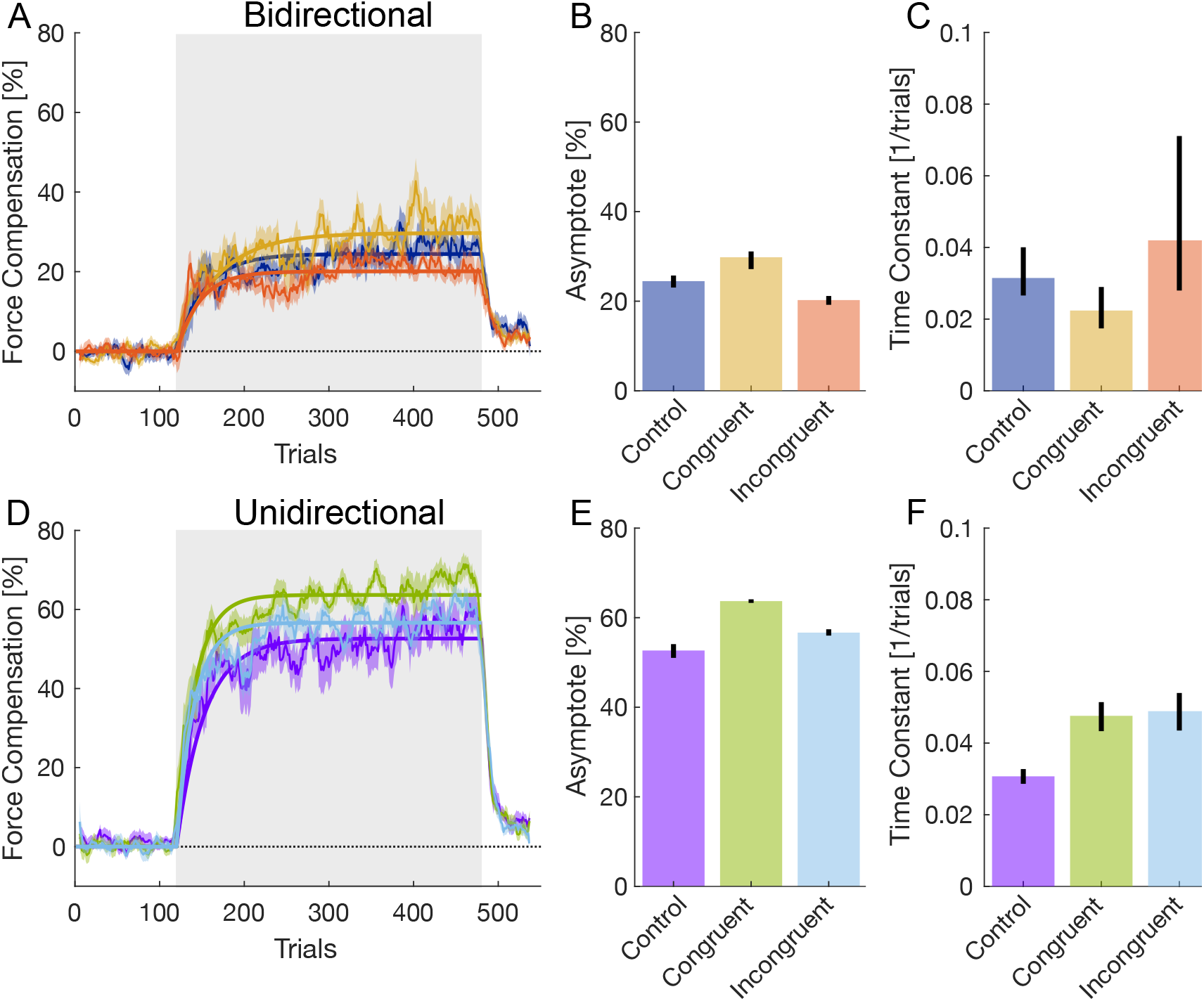
Exponential fit of adaptation. **A.** Bidirectional Force field groups. The best-fit exponential curve during the exposure phase of the experiment (solid line) and the mean and standard error of the mean across trials (thin line and shaded region) is shown. Shaded grey region indicates the period of force field exposure. **B.** The mean best-fit asymptote for each of the three bidirectional force field groups. Error bars represent 95% confidence intervals of the parameters. **C.** The mean best-fit time constant for each of the three bidirectional force field groups. Error bars represent 95% confidence intervals of the parameters. **D.** The best-fit exponential curve during adaptation to the unidirectional Force field groups. **E.** The best-fit asymptote for each of the three unidirectional force field groups. **F.** The best-fit time constant for each of the three unidirectional force field groups.

### Model Results

We simulated the learning process for each of the experimental conditions using state estimation based learning. In this learning scheme (Fig 5A), the state estimator integrates information from multiple sensory modalities, if available, to estimate the scaling factor from noisy observations. Examples for data sets available to the state estimator according to each experimental condition are depicted in Figure 5B. Using these data sets, the state estimator estimates the scaling factor which determines the level of adaptation to the force field. We calculated the difference between estimated and actual scaling factor. Figure 5C shows the learning trends for each condition during adaptation to the unidirectional force field. For this adaptation protocol, the trends predicted by the model are similar to the experimental observed trends (Fig 2 & 3). We found that for the congruent condition, the scaling factor estimation converges faster than the other conditions since in this condition the state estimation can utilize both proprioceptive and visual information together with higher sensitivity to the observation. For the incongruent condition we found that lower observation sensitivity can lower the learning rate. This lower sensitivity is due to our assumption regarding the reliability of the observations during this condition which are opposite to each other. For adaption to the bidirectional force field, we found again a faster adaptation rate for the congruent condition compared with the incongruent and control conditions (Fig 5D). Here, we simulated even lower observation sensitivity for the incongruent condition which resulted in a learning rate which was slower compared with the control condition.

**Figure 5:**
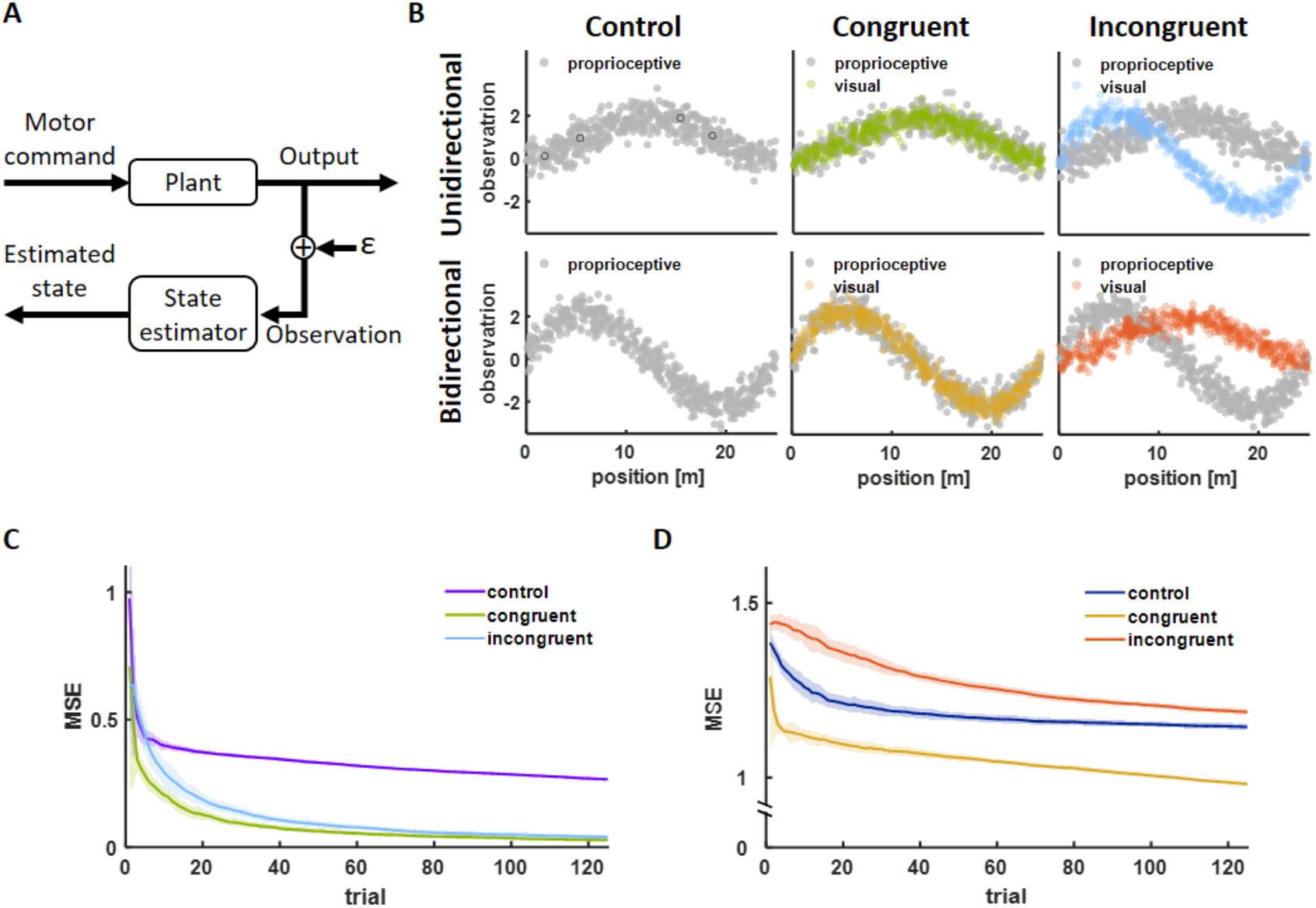
State estimation based adaptation. **A.** Block diagram of state estimation. The motor command sent from the brain changes the state of the arm (plant) which is sensed by multiple sensory modalities such as proprioception or visual. The role of the state estimator is to estimate the state using integration between information sources and reduce the effect of noise (ε) which distort the sensory signals. **B.** Example for simulated sensory information regarding the scaling factor k for each experimental condition across all trials. In each trial we simulated that the system acquires four randomly chosen data points, for example, the four data points marked using black circles in the upper left panel. Top raw of panels represent examples for proprioceptive (grey dots) and visual (green dots for congruent and light blue dots for incongruent conditions) information when the force field is unidirectional. Bottom panels represent examples for proprioceptive (grey dots) and visual (yellow dots for congruent and red dots for incongruent conditions) information when the force field is bidirectional. **C.** Difference between estimated scaling factor, 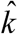, and actual value k measured using mean squared error (MSE) between the two signals for adaptation to the unidirectional force field. For the congruent condition (green line) there is a faster convergence between the true and estimated scaling factor values compared with the incongruent (light blue) and control (purple) which predicts that in this condition participants could estimate faster and more accurate the nature of the force field. For the control condition, the visual nature of the force field is absent which lowers the learning rate. For the incongruent condition, we simulated higher uncertainty in the scaling factor prediction which up-regulated the learning rate compared with the control condition. **D.** Same as C but for adaptation to the bidirectional force field. Here, we simulated lower uncertainty in the scaling factor prediction which down-regulated the learning rate in the incongruent condition (red curve) compared with the congruent (yellow curve) and control (blue curve).

We also found that the ability of the state estimation to accurately estimate the scaling factor of the bidirectional force field was reduced compared with the unidirectional force field. This slower adaptation for the bidirectional force field was also evident in the experimental results (Fig 2 & 3). This difference between force fields was evident even for conditions in which parameters had the same values, for example, in the congruent and control conditions.

## Discussion

In this study, we examined whether additional visual information regarding the nature of external dynamics affects the rate of adaptation to these forces. We provided participants with online visual representation of forces that could accurately or falsely represent the experienced forces and examined if these visual cues assist force field adaptation, especially when the cues were congruent with the experienced forces. Three groups experienced a unidirectional force field, while the other three groups experienced a bidirectional force field. Using these unidirectional or bidirectional velocity-dependent force fields, we have compared different responses of additional visual cues regarding the forces – constant, congruent with the dynamics (cues matched with the type of force field) or incongruent with the dynamics. We observed higher adaptation levels for groups that adapted to the unidirectional force field compared with groups that adapted to the bidirectional force field. For both force field types, the groups in which the visual cues were congruent with the type of force field exhibited faster learning and also higher adaptation level at the end of learning. When the visual feedback was conflicting or not informative, the adaptation rate and final adaptation level were reduced. These results demonstrated that the characteristics of motor learning can change with additional visual information regarding the nature of the environment.

There is contradicting evidence regarding the role of visual feedback in motor planning or adaptation. Congenitally blind individuals are able to adapt rapidly to novel dynamics in the complete absence of visual feedback (DiZio and Lackner 2000). Similarly, studies have shown that people learn both stable and unstable dynamics equally well both with and without visual feedback (Franklin et al. 2007; Scheidt et al. 2005; Tong et al. 2002). However, previous studies have also shown that visual information can alter motor plans and can have a great impact on motor behaviour. For example, visual feedback provides useful information for dynamical control in particular to select different internal models of objects (Gordon et al. 1993). Visual feedback has also been shown to be responsible for learning the direction of the movement and path planning during adaptation (Scheidt et al. 2005). Indeed participants experiencing a visuomotor rotation paradigm are able to successfully perform the task without proprioceptive feedback, leading to the conclusion that the visual signal enables remapping of the planned movement direction (Bernier et al. 2006).

The influence of visual feedback on motor control is not limited to humans; even jumping spiders move their head towards moving objects when their lateral eye captures the motion in order to use their principal eyes to define the event. The retinae of the spider not only provides information about the existence of external stimuli, but also it’s precise orientation, which is translated into a sequence of instructions to orient the spider appropriately (Land 1971). In humans there is some evidence that movements are planned in a manner that suggests the prioritization of visual feedback for certain tasks. For example, Flanagan and Rao (1995) have shown that the human brain will modify the movement path in order to provide a straight path in visual space. Similarly, participants will perform strongly curved movements in order to obtain sufficient visual feedback about the movement (Yeo et al. 2016). However visual feedback does not only take into account motion of the endpoint. It has been shown that our planned trajectory is based on the visual geometric information of the entire system, not just the endpoint, by using a novel mapping between finger movements and the object movement (Danziger and Mussa-Ivaldi 2012). In terms of adapting simultaneously to two opposing force fields, certain types of visual feedback can be used as a contextual cue to separate the learning of these dynamics (Forano et al. 2021; Howard et al. 2020; Howard et al. 2012; Howard et al. 2013; Howard et al. 2015). The fact that only specific types of contextual information allows the learning of two opposing force fields, suggests that only specific signals trigger a switch between or access specific motor memories in a given task.

Compared with most previous studies of learning novel dynamics, in which the visual information provided feedback about motion errors due to the force field or cued that a different force field would be applied, in this study we examined how additional visual information regarding the nature of the force field can affect adaptation. In this case, participants could potentially ignore the added information, especially for the incongruent visual feedback condition, and have similar adaptation as with no additional visual feedback. Despite this, we found that the visual information affected adaptation, suggesting an inability to neglect the visual representation of forces. This is in line with other results of manipulating objects while experiencing incongruent visual feedback. For example, incongruent visual information regarding the tip of an inverted pendulum decreased the ability of participants to stabilize the pendulum despite having reliable haptic information regarding the mechanical properties of the pendulum (Česonis et al. 2019; Franklin et al. 2018). The inability to discard the incongruent visual information suggests involvement of unconscious learning processes in which this visual information can enhance or reduce the learning rate.

Indeed, several studies suggested that visual feedback triggers a more conscious, explicit strategy of action when adapting to external dynamics. That is, the deviation of the visual representation of the hand, usually a cursor, from the desired straight line movement allowed participants to identify the force nature, such as the force direction, and aim or move differently. For example, Hwang et al. (2006) showed that visual information about hand position but not proprioceptive information can make participants aware of the presence of a force field. They showed that this awareness caused an improvement in learning for both cases where visual information was reliable and when it was not. However, this study examined visual information which was directly linked with the motion error due to forces. In a recent study Zhou and colleagues (2022) contrasted motor learning performance between two experimental groups in a reach adaptation task in order to assess the extent to which explicit visual feedback facilitates motor learning. Specifically, one group of participants experienced a velocity dependent force field (i.e., received both visual and proprioceptive feedback), whereas the other group moved in force channels and was provided visual feedback concerning the lateral force they produced as well as the force required overcome the force field (i.e., plotted the velocity-dependent force field curve). They found that decay was faster in the group that only received explicit visual feedback, although the single-trial recall was similar between the two groups. The authors concluded that motor adaptation can rely on explicit visual feedback, although adaptation in this case is less stable than that based on experiencing multisensory errors following physical perturbations. While this study provided online visual feedback of the forces during the movement, the visual traces remained on the screen until after the end of the movement. This is slightly different than previous work which provided a visual picture of the forces prior to the start of the movement (Osu et al. 2004). Both of these are quite different to the approach of our study in which only the instantaneous visual feedback was connected to the dynamics of the environment. As we know that continuous and terminal feedback effects learning in a very different manner (Heuer and Hegele 2008), it is possible that all of these studies produce changes in adaptation through different mechanisms.

To explain the experimental results, we proposed a computational model that predicts the pattern of adaptation based on a state estimation. In this model, the state estimator forms a representation of the external force field by integrating proprioceptive information with additional visual information that represents the pattern of external forces. The state estimator receives information from the two modalities and updates the representation of the environment as more information is accumulated. Our model suggests that the learning rate and the adaptation steady state value depend on the uncertainty levels we have of the external dynamics representation. That is, when the two sources of information are in agreement, such as in the congruent condition, the state estimator is more uncertain in the current representation and thus change more rapidly the estimation based on new observations. This is in line with other state estimation based adaptation (Burge et al. 2008; Wei and Koerding 2010) in which the learning rate was altered with changes in the noise in the sensory feedback. Here however, since we did not alter sensory noise but provided multiple sensory inputs with different levels of congruency between them we suggest a complementary way of controlling adaptation rate which also supports the idea of a Kalman filter (Kalman 1960) as a way of implementing state estimator based adaptation.

A second assumption of the suggested state estimation adaptation model is the difference in uncertainty levels according to the complexity of the force field. For the incongruent condition, we assumed higher ability of the model to modify the representation of the scaling factor for the unidirectional force field compared with the bidirectional force field. This allowed different adaptation pattern in the incongruent condition between these two types of force fields. We suggest that in less complex force fields, such as the unidirectional field, participants were able to better learn the force pattern despite the incongruent additional visual information. Similar to increased generalization ability of low complexity force fields (Thoroughman and Taylor 2005) or higher adaptation rate when visual cues are less variable (Howard et al. 2017), we suggest that the complexity of the unidirectional force pattern was low enough so to have different adaptation nature such as the ability to disregard or depress the effect of the additional visual information. For the congruent and control conditions, the model predicted different levels of estimation despite setting the uncertainty level to have the same value between unidirectional and bidirectional fields. This was consistent with the experimental results and suggests that in general the bidirectional force field was more difficult to estimate and that it might require much more extensive training in order to achieve the same level of adaptation.

When people are exposed to a fully or partially unknown environment, any information about the characteristics of the environmental dynamics is valuable. Both visual and proprioceptive feedback can be used both to estimate the current state of the body and information about how to correct the movements to minimize the error. Here we have shown that even additional visual signals can be used to increase adaptation to novel dynamics when such information relates to the dynamics of the environment. In this study it is unclear whether such an effect was driven by changes in explicit or implicit learning. However, the computational model also replicated the experimental differences across conditions. This model suggested that the major difference was the use of this additional visual information to improve state estimates during adaptation. While this is only one possible explanation, out of many, for the results, it would suggest a more implicit explanation for the effect of congruent visual cues on the speed of adaptation. This matches with recent work suggesting a strong component of implicit learning in force field adaptation (Forano et al. 2021; Schween et al. 2020). Further studies would be needed to confirm whether these visual cues are used within state estimation as we suggest with our model. One possible test of this would be to investigate the role of these online visual cues in dual adaptation paradigms (Howard et al. 2013) as we have suggested that strength of contextual cues depends on their use within state estimation (Forano et al. 2021; Howard and Franklin 2016; 2015; Howard et al. 2020; Howard et al. 2012). While we can only speculate on the mechanism which drives this increase in adaptation, it is clear that congruent visual cues increase the adaptation to novel dynamics compared with either incongruent or the absence of visual cues. Such effects might be useful for rehabilitation or training, especially if they can contribute through improved state estimation.

## Acknowledgment

We thank Yang Wang with her input on the experimental design, and Kevin S.H. Teo and Vijay Maharajan for their help with the experimental recordings.

## Author Contributions

Experimental Design: DWF

Running experiments: SF and DWF

Analysis: DWF and MD

Model Simulation: RL

Wrote Manuscript: SF, RL and DWF

Edited Manuscript: SF, RL, MD and DWF

